# Mechanism of BceAB-type transporter: Resistance by lipid II flipping

**DOI:** 10.1101/2020.03.05.978734

**Authors:** Julia Zaschke-Kriesche, Sandra Unsleber, Irina Voitsekhovskaia, Andreas Kulik, Lara V. Behrmann, Nina Overbeck, Kai Stühler, Evi Stegmann, Sander H.J. Smits

## Abstract

Treatment of bacterial infections are the great challenge of our era due to the evolved resistance mechanisms against antibiotics. The Achilles heel of bacteria is the cell wall especially during the needs of its synthesis and cell division. Here lipid II is an essential cell wall precursor component synthesized in the cytosol and flipped into the outer leaflet of the membrane prior to its incorporation into the cell wall.

Compounds targeting the cell wall or its biosynthesis precursors have been around for decades and have been used as antibiotics against bacterial infections like meningitis, pneumonia and endocarditis. Antimicrobial peptides (AMPs) have proven to be a promising weapon against multiresistant bacteria. However, the <u>**B**</u>a<u>**c**</u>itracin <u>**e**</u>fflux (BceAB)-type ATP binding cassette transporters expressed in the membrane of human pathogenic bacteria have been shown to confer resistance to these alternative antibiotics, thereby hampering their medical development.

In *Streptococcus agalactiae* COH1 the BceAB-type transporter NsrFP (*Sa*NsrFP) confers high-level resistance against the antimicrobial peptide nisin, a member of the lantibiotic subfamily. We showed that *Sa*NsrFP provides a novel resistance mechanism by flipping lipid II back into the cytosol, thereby preventing the binding of nisin as well as other lipid II targeting compounds. This is intriguing since a relatively simple reaction mediates resistance to human pathogenic bacteria to lipid II targeting antibiotics, regardless of their structure.

**Significance Statement:** The ABC-transporter NsrFP from *Streptococcus agalactiae* (*Sa*NsrFP) belongs to the BceAB-type transporters. Several BceAB-type transporters are known to confer resistance against multiple antimicrobial peptides. In this study a new resistance mechanism was identified, which is based on the reduction of the number of cell wall precursor lipid II molecules on the cell surface mediated by *Sa*NsrFP. *Sa*NsrFP flips lipid II, which are considered to be the target for many antibiotics, back into the cytoplasm. With this newly gained knowledge about the resistance mechanism of BceAB-type transporters, novel strategies can be established to overcome or bypass this resistance in human pathogenic bacteria.

## Introduction

Bacterial infection cause over 150,000 death every year and are the major threat for humans (1). The treatment of many infections is possible through the development of antibiotics, which exist since 1929, starting with penicillin (2). In recent years however antibiotic resistance has become a great challenge, and targeted bacteria evolved several different resistance mechanisms (3). The Achilles’ heel of bacteria was shown to be biosynthesis of the cell wall and here especially the essential cell wall precursor lipid II (4). The biosynthesis of lipid II, it’s incorporation into the nascent cell wall as well as the recycling as well as the recycling pathway of undecaprenylpyrophosphate (UPP), have been ideal targets for novel antimicrobial compounds (5). The bacterial cell wall is composed of peptidoglycan (PGN), a polymer consisting of repeating units of *N*-acetylmuramic acid (MurNAc) and *N*-acetylglucosamine (GlcNAc). Attached to the N-acetylmuramic acid is a peptide chain consisting of three to five amino acids. The peptide chain is cross-linked to the peptide chain of another strand forming the stable network(6). The biosynthesis of the PGN requires several steps, which are evolutionary conserved in all bacteria species (7). Briefly, lipid II is composed of an undecaprenyl-pyrophosphate (UPP) anchor, the two amino sugars GlcNAc and MurNAc and a covalently attached pentapeptide, is built up in the cytoplasm via several consecutive enzymatic reactions and afterwards flipped to the extracellular space (or periplasm for Gram-negative bacteria), still anchored to the membrane *via* UPP (8). Subsequently, the GlcNAc-MurNAc-pentapeptide subunit is incorporated into the nascent PGN, leaving UPP residing in the membrane. UPP is subsequently dephosphorylated to undecaprenyl phosphate (UP), which is flipped back into the cytoplasm, and implemented into a new PGN synthesis cycle (9) (Figure 1). Interruption of this biosynthetic pathway at any stage effectively results in inhibition of bacterial growth, leading to turgor pressure and consequently to cell death (10). In cases where lipid II is bound by antibiotics or when the enzymes that catalyze its incorporation into PGN are inactivated by the antibiotics, the lipid carrier molecule UP cannot be recycled and thus conclusively no new lipid II can be synthesized. This leads to the growth inhibition described above.

**Figure 1:**
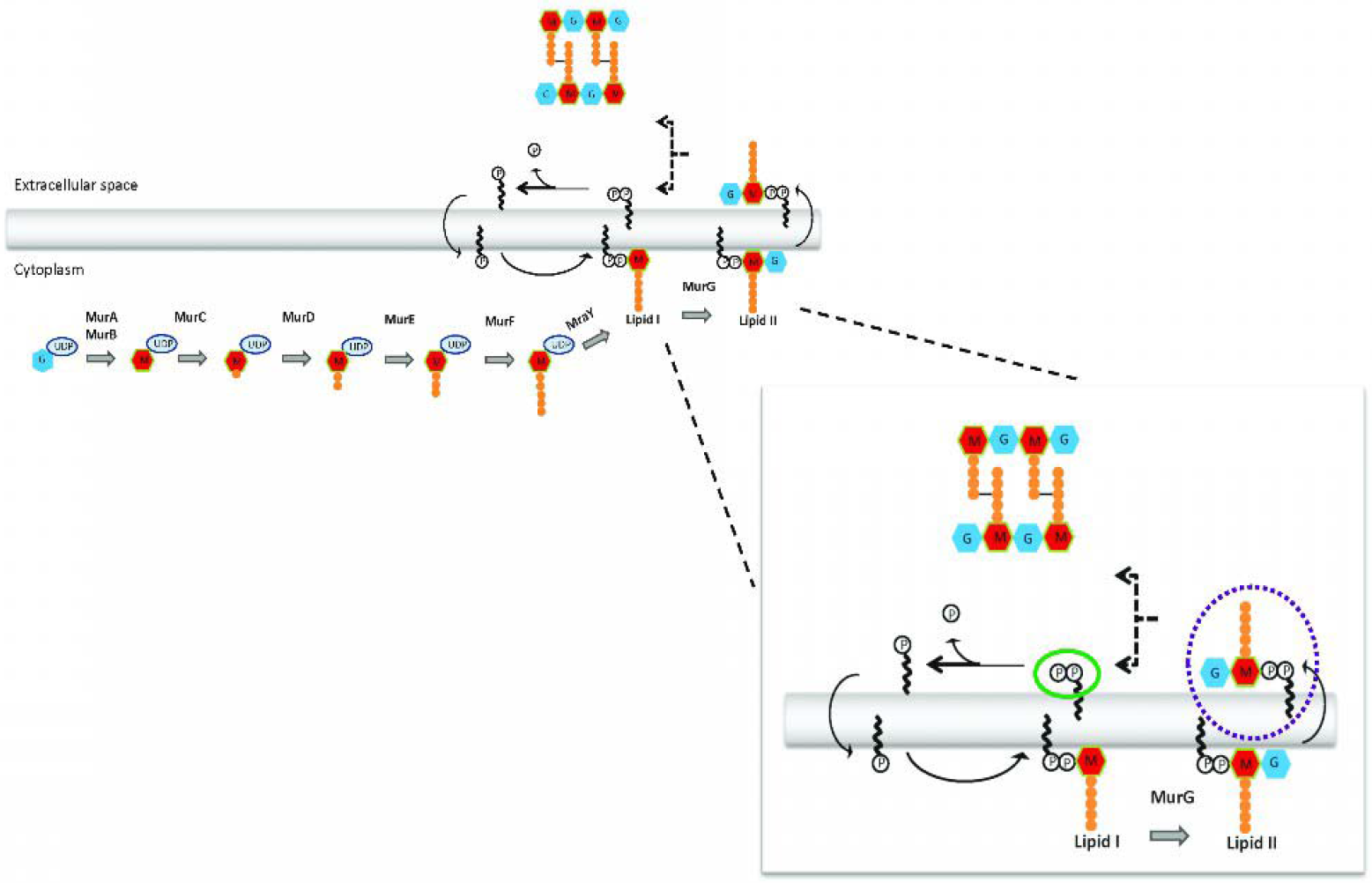
Schematic view of peptidoglycan synthesis. Phosphates are marked with an P, undecaprenyl as a black curved line, uridine phosphate (UDP) in light blue, GlcNAc in blue, MurNAc in red and amino acids of the pentapeptide in orange. Enlarged step of the peptidoglycan synthesis showing targets of bacitracin, undecaprenyl pyrophosphate (green circle) and target of antimicrobial peptides like nisin, gallidermin, lysobactin, ramoplanin and vancomycin, lipid II (violette, dotted circle).

Antibiotics bind to various moieties of lipid II for example the pyrophosphate moiety (lantibiotics like nisin and gallidermin) (11–13) or the pentapeptide moiety (glycopeptides like vancomycin) (14–16) (Figure 1). are small antimicrobial peptides (AMPs) that bind specifically to lipid II (11), such as the lipoglycodepsipeptide ramoplanin and the acylcyclodepsipeptide lysobactin A particular class are the small antimicrobial peptides (AMPs), that bind specifically binding to lipid II (11), like the lipoglycodepsipeptide ramoplanin and the acylcyclodepsipeptide lysobactin (5, 17–19).

In addition to antibiotics specifically binding lipid II, there are other antibiotics that inhibit PGN biosynthesis in different phases: for example the cyclic peptide bacitracin, which binds to the undecaprenyl-pyrophosphate (UPP), thereby preventing the back flipping of the lipid carrier into the inside of the cell consequentially resulting into the accumulation of lipid II molecules into the cytoplasm and to the interruption of PGN biosynthesis (20) (Figure 1).

A rather novel family of ABC-transporters, the <u>**B**</u>a<u>**c**</u>itracin <u>**e**</u>fflux (Bce) type transporters have been identified to confer high-level resistance against bacitracin as well as against lantibiotics like nisin and gallidermin in *Bacillus subtilis, Staphylococcus aureus* and *Streptococcus agalactiae,* respectively (21). Genome analysis revealed the presence of these transporter especially in human pathogenic bacteria (22). The BceAB-type transporter, which was first discovered in *B. subtilis* (23) is encoded on one operon together with the BceRS-like two component system (TCS) which regulates the expression of the transporter (24). Several putative mechanisms have been proposed for BceAB-type transporter ranging from antimicrobial peptide removal from the membrane (25), operating as an exporter (26) or flipping UPP, a sub-product of the peptidoglycan synthesis (27). More recently a study proposed a target-AMP dissociative mechanism for BceAB-type transporters (28).

We aimed to characterize the resistance mechanism of NsrFP from the human pathogenic bacteria *S. agalactiae*, which has been shown to confer resistance to lantibiotics, by combining different techniques such as cell growth assays, cell wall precursor analysis and whole cell proteome analysis. Surprisingly, we observed a novel resistance mechanism by which lipid II is expected to fold back into the cytosol, thereby preventing antibiotics from binding to the target. This mechanism is actually very sophisticated because it provides the bacterial cell resistance against a large number of structurally unrelated compounds.

## Results

### *Sa*NsrFP confers high resistance against bacitracin

NsrFP from the human pathogen *Streptococcus agalactiae* COH1, *Sa*NsrFP has been shown to confer resistance against the lantibiotic nisin, and structurally related compounds like the natural variant nisin H and gallidermin by recognizing and binding to the N-terminus of these lantibiotics (29). We extended this resistance spectrum studies to a structurally divers, rather unrelated group of antibiotics, including ramoplanin, vancomycin and lysobactin binding at different parts of lipid II (30), as well as bacitracin, which exclusively binds to the lipid carrier UPP (20) by comparison of the IC_50_ of a sensitive *L. lactis* strain as well as a strain expressing *Sa*NsrFP (*L. lactis* NZ9000NsrFP) or the inactive mutant *Sa*NsrF_H202A_P (*L. lactis* NZ9000NsrF_H202A_P) (26).

Compared to *L. lactis* wildtype (WT), *L. lactis* producing *Sa*NsrFP mediated a moderate resistance against lipid II binders (2-6-fold), however a striking high-level resistance against bacitracin (350-fold)(see Figure 2A, SI Figure 1 and Table 1). *L. lactis* producing the mutated variant of the ABC-transporter *Sa*NsrF_H202A_P displayed similar IC_50_ values against the applied antibiotics as *L. lactis* WT indicating that *Sa*NsrFP is actively conferring the displayed resistance. Western blot analysis of the membrane fraction revealed that both, *Sa*NsrFP and the dead mutant variant *Sa*NsrF_H202A_P were produced at the same level (Figure 2B), whereas the control cells of *L. lactis* WT containing the empty vector pIL-SVCm (NZ9000Cm) (*L. lactis* NZ9000Cm) showed no *Sa*NsrFP production.

**Figure 2:**
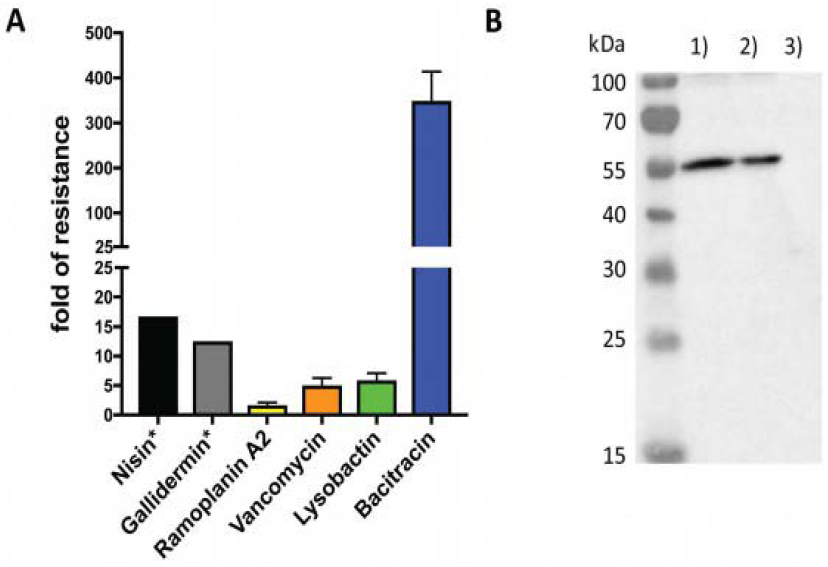
**A) Fold of resistance of *L. lactis* NZ9000NsrFP against *L. lactis* NZ9000Cm** calculated with determined IC_50_ of ramoplanin A2 (yellow), vancomycin (orange), lysobactin (green) and bacitracin (blue). Values for nisin and gallidermin were taken from Reiners et al. (2017) (26) and marked with an asterisk. Values were calculated from at least 5 independent measurements. **B) Production of *Sa*NsrFP (1) and *Sa*NsrF_H202A_P (2)** in *L. lactis* NZ9000Cm as well as the empty vector pIL-SV in *L. lactis* NZ9000Cm monitored *via* western blot with a polyclonal antibody against the extracellular domain of *Sa*NsrP.

Since *Sa*NsrFP confers resistance against structurally unrelated compounds, we concluded that *Sa*NsrFP is neither able to inactivate such large number of different compounds nor simply binding them but rather confers resistance by interfering with cell wall biosynthesis components like lipid II, UPP or UP, present at the bacterial membrane exterior. The high-level resistance observed for the UPP binder bacitracin, suggested that *Sa*NsrFP interferes directly with lipid II or the recycling of UPP (Fig. 1 and 2A).

### *Sa*NsrFP enables normal growth in the presence of bacitracin

In order to investigate whether *Sa*NsrFP interferes with PGN synthesis we analysed the influence of the production of *Sa*NsrFP on the growth of *L. lactis.* The growth of the three strains *L. lactis* NZ9000Cm, *L. lactis* NZ9000NsrFP and *L. lactis* NZ9000NsrF_H202A_P was monitored online over a time period of 500 minutes (Figure 3A). A sublethal concentration of 0.3 nM nisin was added to the cultures at an OD of 0.5 (31 37)) to induce the expression of the *Sa*nsrfp gene. *L. lactis* NZ9000NsrFP entered the stationary phase two hours later than *L. lactis* NZ9000NsrF_H202A_P and *L. lactis* NZ9000Cm strain (Figure 3A).

**Figure 3:**
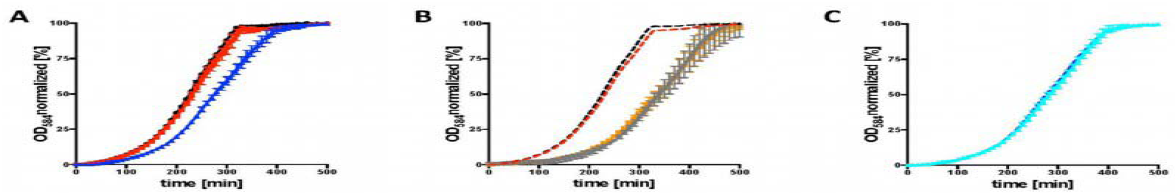
**A) Growth curve** of *L. lactis* NZ9000NsrFP (blue), *L. lactis* NZ9000NsrF_H202A_P (red) and *L. lactis* NZ9000Cm (black) induced with 0.3 nM nisin. **B) Growth curve** of *L. lactis* NZ9000NsrF_H202A_P (orange) and *L. lactis* NZ9000Cm (grey) induced with 0.3 nM nisin and additional 100 nM bacitracin compared to *L. lactis* NZ9000NsrF_H202A_P (red) and *L. lactis* NZ9000Cm (black) without bacitracin. **C) Growth curve** of *L. lactis* NZ9000NsrFP (blue) induced with 0.3 nM nisin and *L. lactis* NZ9000NsrFP (turquoise) with additional 100 nM bacitracin.

We repeated these experiments supplemented with an additional 100 nM bacitracin. The use of a sublethal concentration of bacitracin (100 nM) causes significantly slower cell growth of *L. lactis* NZ9000Cm and NZ9000NsrF_H202A_P (Figure 3B grey and orange). This retardation of growth upon addition of bacitracin is known for many bacterial cells, for example methicillin resistant *S. aureus* and Group B *Streptococcus* (32) and is caused by the binding of bacitracin to UPP (20).

Intriguingly, *L. lactis* NZ9000NsrFP did not show reduced growth (Figure 3C) in the presence of bacitracin. This implicated that the *L. lactis* NZ9000NsrFP is resistant to bacitracin and that this resistance is linked to the fact that the bacitracin target UPP is not accessible in this strain. However, since the cells are still able to grow, albeit at a slower rate, lipid II synthesis does not appear to be completely inhibited.

### SaNsrFP prevents accumulation of peptidoglycan precursors after addition of bacitracin

Bacitracin blocks cell wall formation by interfering with the dephosphorylation of UPP and preventing the formation of UP. As the carrier of lipid II, however, UP is essential for PGN synthesis; when lipid II is incorporated into the growing PGN chain, UPP is released. Before a new molecule of lipid II can be linked to the carrier, it has to be dephosphorylated to UP. As a consequence, the inhibition of UP regeneration leads to the accumulation of intracellular PGN precursors (33 40)).

To further address the mode of action of *Sa*NsrFP we analysed the PGN precursors accumulated in the cytoplasms of *L. lactis* NZ9000NsrFP, *L. lactis* NZ9000Cm *and L. lactis* NZ9000NsrF_H202A_P grown in the presence of bacitracin. HPLC/MS analysis revealed the presence of the characteristic PGN precursors UDP-MurNAc-Ala-Glu-Lys-Ala-Ala (1148.4 m/z^-1^) and UDP-MurNAc-Ala-Glu-Lys-(Asn)-Ala-Ala (1263.4 m/z^-1^) in *L. lactis* NZ9000Cm (Figure 4).

**Figure 4:**
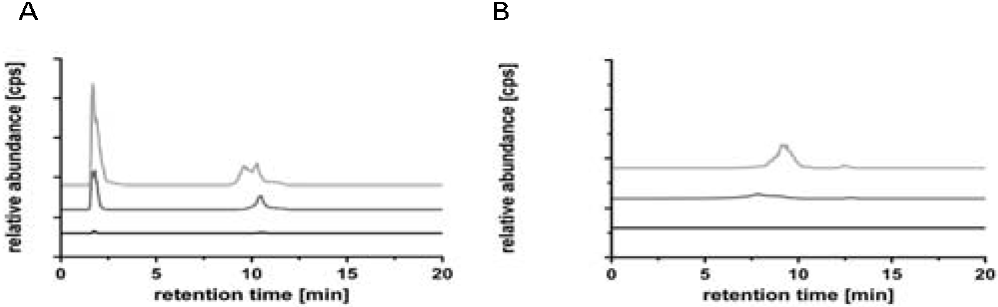
Relative abundance of obtained mass in cps against retention time in min of precursor accumulation after treatment with bacitracin. **A)** Extracted ion Chromatograms (EIC) spectrum for UDP-MurNAc-Ala-Glu-Lys-Ala-Ala (1148.4 m/z^-1^) and **B)** EIC spectrum for UDP-MurNAc-Ala-Glu-Lys(Asp)-Ala-Ala (1263.4 m/z^-1^) of the strains NZ9000Cm (light grey), NZ9000NsrF_H202A_P (grey) and NZ9000NsrFP (black).

Interestingly, no accumulation of PGN precursors was observed in *L. lactis* NZ9000NsrFP. These findings, together with the results obtained in the resistance test, suggest that *Sa*NsrFP prevents the binding of bacitracin to UPP and thus the inhibition of PGN synthesis. This hypothesis is further supported by the results obtained for *L. lactis* NZ9000NsrF_H202A_P. Here, only a slightly reduced PGN precursor accumulation was observed compared to the bacitracin-sensitive *L. lactis* NZ9000Cm control strain (Figure 4A).

Since bacitracin did not inhibit the PGN synthesis in *L. lactis* NZ9000NsrFP, two different mechanisms of action have been postulated for *Sa*NsrFP: either *Sa*NsrFP flips lipid II itself or UPP back into the cytoplasm or *Sa*NsrFP inhibits the binding of bacitracin to UPP as recently postulated (28).

### *Sa*NsrFP causes downregulation of the proteins involved in peptidoglycan synthesis

We analysed the whole proteome of NZ9000NsrFP, NZ9000Cm and NZ9000NsrF_H202A_P *L. lactis* strains grown under identical conditions (see Material and methods) by mass spectrometry. The analyses of the proteomes leaded to the identification of 894 proteins (identified by at least two unique peptides in each strain). The comparison between *L. lactis* NZ9000Cm and NZ9000NsrFP revealed 315 differentially expressed proteins (Figure 5A). 339 proteins showed differential abundances between the *L. lactis* strains NZ9000NsrF_H202A_P and NZ9000NsrFP (Figure 5B). This high number of differentially produced proteins implicated that *L. lactis* NZ9000NsrFP has to respond significantly to counteract the effects mediated by the expression and presence of solely the *Sa*NsrFP transporter.

**Figure 5:**
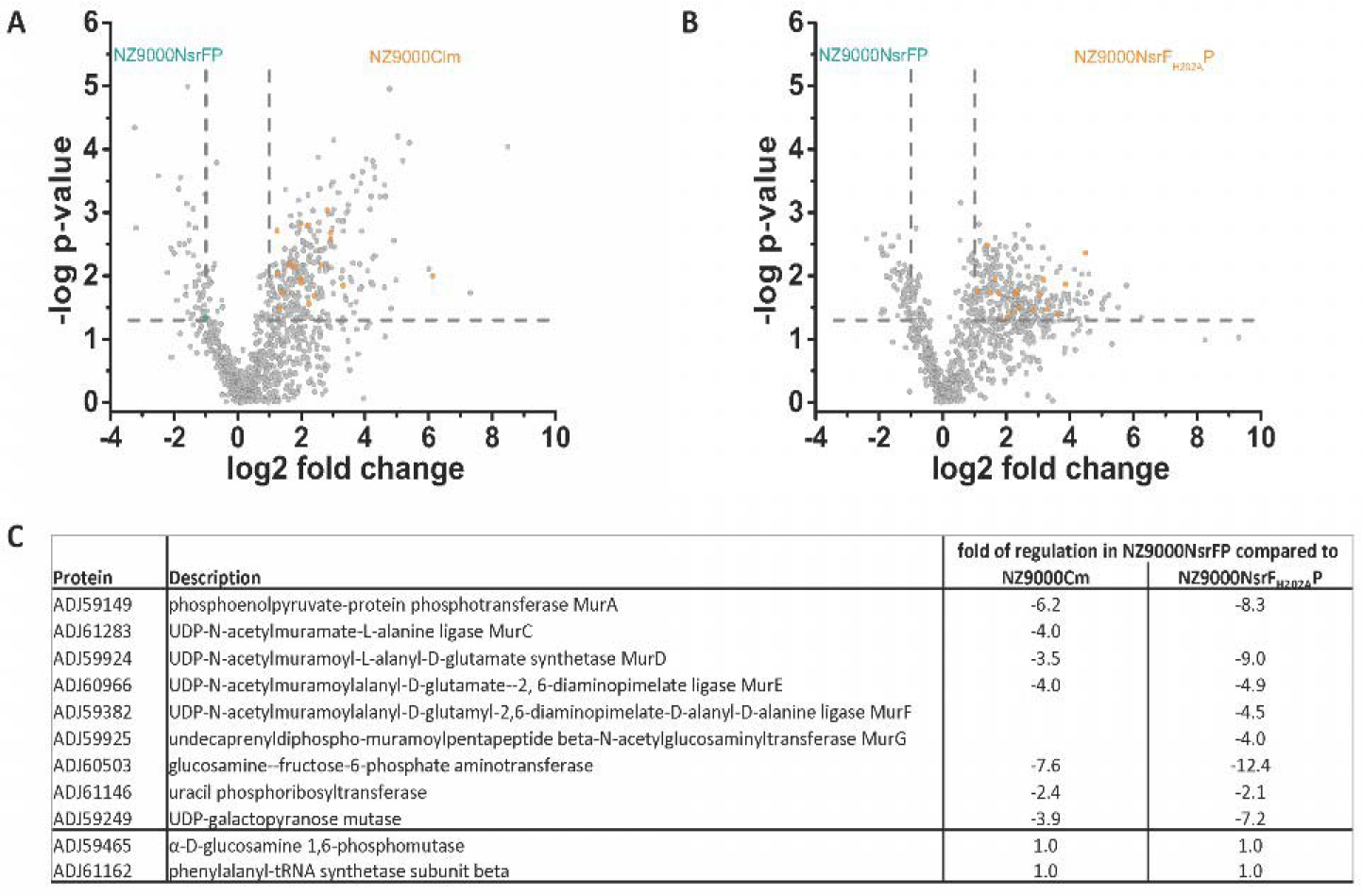
**A) Volcano plot** of the proteome analysis of **NZ9000NsrFP against NZ9000Cm** and **B) NZ9000NsrFP against NZ9000NsrF_H202A_P.** Proteins involved in the cell wall synthesis are highlighted in orange if upregulated in NZ9000Cm (A) and NZ9000NsrF_H202A_P (B) as well as highlighted in blue if upregulated in NZ9000NsrFP. **C) Selected proteins** of the proteome analysis with their description and their fold of down regulation in NZ9000NsrFP compared to NZ9000Cm and NZ9000NsrF_H202A_P.

In depth analysis displayed that in particular the production of proteins involved in cell wall synthesis is reduced in *L. lactis* NZ9000NsrFP (Figure 4A & B and SI Figure 3). Among them the phosphoenolpyruvate-phosphotransferase MurA (ADJ59149), is produced 6.2 fold less in the *L. lactis* NZ9000NsrFP compared to *L. lactis* NZ9000Cm; UDP-*N*-acetylmuramate-L-alanine ligase MurC (ADJ61283), 4.0 fold, UDP-*N*-acetylmuramoyl-L-alanyl-D-glutamate synthetase MurD (ADJ59924) the 3.5 fold, UDP-*N*-acetylmuramoylalanyl-D-glutamate-2, 6-diaminopimelate ligase MurE (ADJ60966), 4.0 fold less produced in the control strain as well as UDP-*N*-acetylmuramoylalanyl-D-glutamyl-2,6-diaminopimelate-D-alanyl-D-alanine ligase MurF (ADJ59382) and undecaprenyldiphospho-muramoylpentapeptide beta-*N*-acetylglucosaminyltransferase MurG (ADJ59925), which are reduced 4.6 fold and 4.0 fold compared to *L. lactis* NZ9000NsrF_H202A_P, respectively (Figure 5C). Further several proteins involved in the synthesis of individual components of the lipid II synthesis, like uracil phosphoribosyltransferase or the glucosamine-fructose-6-phosphate aminotransferase, responsible for UMP and glucosamine-6phosphate synthesis, respectively, also showed a reduced production in *L. lactis* NZ9000NsrFP as well (Figure 5C). Further several proteins involved in the synthesis of single components of the lipid II synthesis, like uracil phosphoribosyltransferase or the glucosamine-fructose-6-phosphate aminotransferase, responsible for UMP and glucosamine-6phosphate synthesis, respectively, are down regulated in NZ9000NsrFP as well (Figure 5C and SI Figure 3).

To verify whether the differences in protein production are indeed related to the presence of the ABC-transporter *Sa*NsrFP we also examined proteins of other metabolic pathways, such as amino sugar metabolism or translation, where the protein α-D-glucosamine-1,6-phosphomutase or phenylalanyl-tRNA synthetase beta subunit, respectively, showed no differences in protein production among all three tested strains This proves that the observed differences were indeed due to the presence of *Sa*NsrFP.

The reduced production of the key enzymes of the lipid II cycle is remarkable and suggests that the biosynthesis of new lipid II molecules occurs with less efficiency in the *L. lactis* NZ9000NsrFP strain. This corresponds to the reduced growth behavior observed in the growth analysis (see above) and proofs that lipid II is indeed the actual substrate of *Sa*NsrFP.

## Discussion

Previous studies showed that the BceAB-type ABC transporter *Sa*NsrFP confers resistance against the lantibiotic nisin and gallidermin (26), both binding to the pyrophosphate moiety of the cell wall precursor lipid II (34). Therefore, in this study we addressed the substrate specificity in a broader context. We observed that *Sa*NsrFP confers high-level resistance against the cyclopeptide bacitracin, which binds to UPP (20) with an exceptional high fold of resistance of 349.15 ± 64.9 (SI Table 1). Several other characterized BceAB-type transporters have been characterized also conferring a high resistance against bacitracin. Interestingly also the AnrAB transporter from *Listeria monocytogenes* (35), VraDE of *S. aureus* (36) as well as the ABC-transporter BceAB of *B. subtilis* display high resistance against bacitracin (37). Like SaNsrFP, however, these transporters exhibit in addition a certain resistance to nisin and gallidermin, suggesting a general mechanism of resistance rather than a resistance specifically against one type of antibiotics.

The binding of bacitracin to UPP or UP normally results in the accumulation of the lipid II precursors within the cytosol. The lack of this accumulation in the *Sa*NsrFP producing cells suggest that in this case bacitracin is unable to bind to its membrane localized target, and the strain thus exhibits high resistance to bacitracin. Together with the observation that the cell growth in the presence of bacitracin is similar to the cells without *Sa*NsrFP suggested that UPP is significantly reduced in the membrane of *SansrFP* expressing cells. This indicates that *Sa*NsrFP actively transports UPP back into the cell thereby reducing the concentration of the bacitracin target at the membrane, and preventing the inhibition of a new lipid II biosynthesis cycle.

This would indicate that *Sa*NsrFP is transporting UPP back into the cell thereby reducing the concentration of the bacitracin target in the membrane, allowing a new lipid II biosynthesis cycle to occur involving the synthesis of the building blocks GlucNAc, MurNAc and pentapeptide. The synthesis of lipid II occurs in the cytoplasm and involves the synthesis of the building blocks GlucNAc, MurNAc and the pentapeptide chain. This requires the enzymatic activity of MurA, MurB, MurC, MurD, MurE, MurF, MraY and MurG, for which we however observed a decreased concentration in the SansrFP expressing cells.

Therefore, we postulated a new SaNsrFP-based resistance mechanism against antibiotics targeting lipid II: SaNsrFP flips lipid II back into the cytosol. As a consequence, less lipid II has to be synthesized *de novo*, thus downregulating the transcription of the lipid II biosynthesis genes. As a result, only a limited amount of lipid II (1.5 x 10^5^ lipid II molecules per cell (38)) is available for cell division and cell growth, explaining the significantly reduced growth of *SansrFP*-expressing cells compared to the control cells. A lower amount of lipid II molecules is further hampering the building of new PGNs (43).

Recent studies revealed that the promotor of *bceA* is ten times stronger when the gene for the UPP phosphatase is knocked out in the bacterial strain (28). The authors assumed that increased UPP levels, corresponding to the absence of UPP phosphatase, trigger useless ATP hydrolysis. Based on our studies depicted here, we conclude that the production of the Bce*AB*-type transporter is increased in order to ensure the flipping back lipid II into the cytoplasm and the reduction of UPP molecules in the extracellular space.

A study on the BceAB transporter of *B. subtilis* postulates flipping of UPP in order to remove the bacitracin target and thus to confer resistance against bacitracin (27). However, UPP flipping would not explain the additional resistance against nisin and gallidermin (37). Therefore, lipid II flipping appears more likely as it also explains the resistance to structurally unrelated antibiotics. BceAB transporters relocate the PGN precursor lipid II and thereby suppress the formation of UPP, which subsequently results in the bacitracin resistance (Figure 2A).

The postulated mechanism of *Sa*NsrFP is demonstrated in Figure 6. Based on all results from this study a flippase activity of the transporter is very likely and will results in resistance against compounds targeting the lipid II cycle. By removing the target molecule lipid II from the extracellular space, less compound e.g. nisin is able to bind and more nisin can be detected in the supernatant, which also explains previous suggestions of an exporting mechanism and further assumptions of the removal from AMPs from the membrane (39). Moreover also an AMP import transport mechanism for BceAB-type transporter is unlikely since it would not declare the results gained in this study or in previous studies (40). A recent study demonstrated that neither import nor inactivation of the substrate are possible mechanisms for BceAB-type transporter (28). Here the authors propose a shielding mechanism of the transporter between the target (e.g. lipid II or UPP) and the AMP based on previous studies and transporter activity studies in presence of more UPP or in presence of C35 isoprenoid heptaprenyl diphosphate (HPP) (28). Those results disprove previous assumptions of an UPP flippase mechanism (27) and indicate a potential target-AMP-dissociation mechanism until the results of this study are taken into consideration.

**Figure 6:**
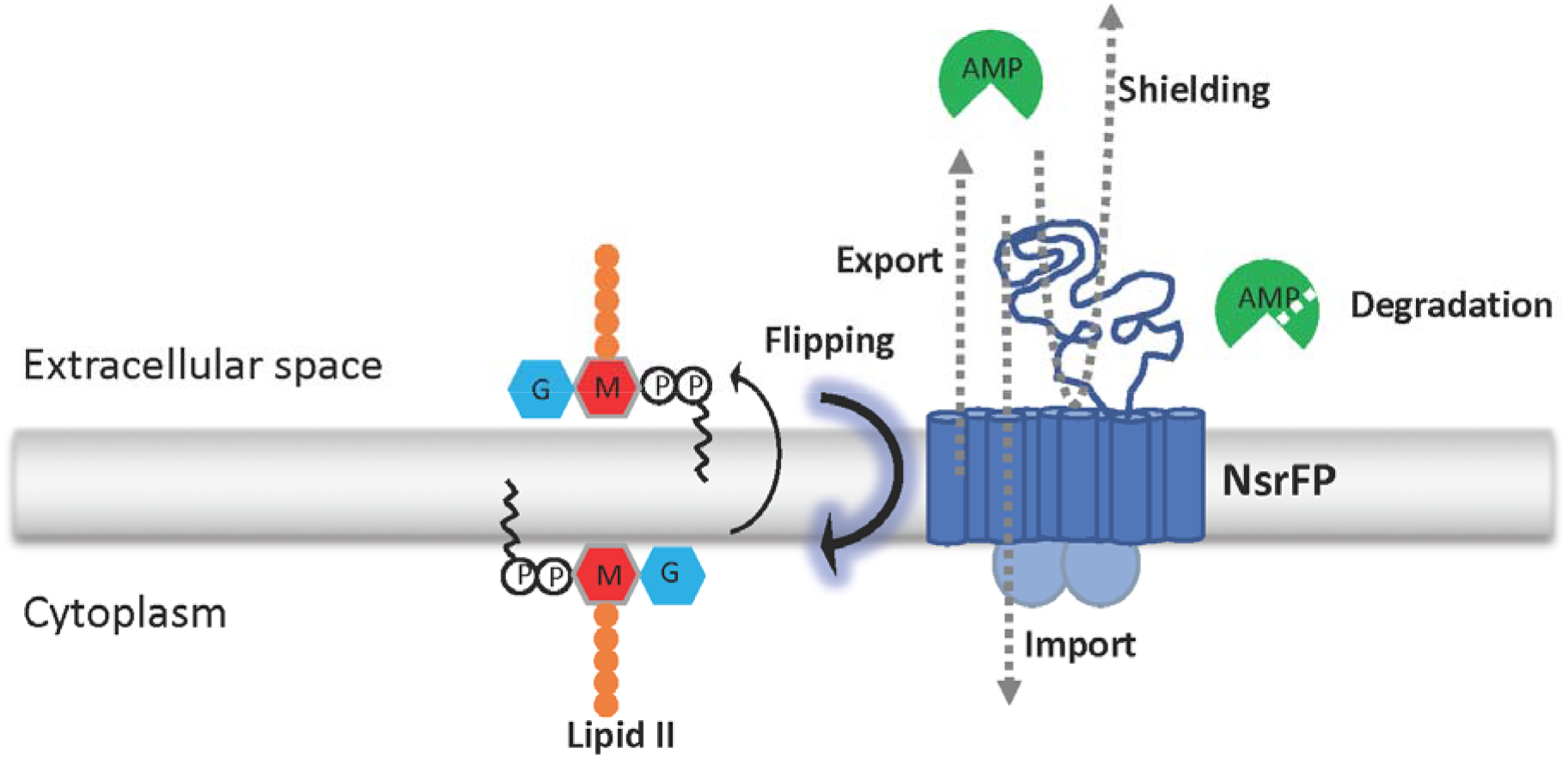
Postulated mechanism of *Sa*NsrFP. Schematic view of lipid II and *Sa*NsrFP. Phosphates are marked with an P, undecaprenyl as a black curved line, GlcNAc (G) in light blue, MurNAc (M) in red and amino acids of the pentapeptide in orange. The transporter *Sa*NsrFP is demonstrated in blue with its large extracellular domain. Dashed grey arrows for possible export, import and shielding mechanism of AMPs (Green) as well as an degradation of AMP (Dashed white line) and a blue highlighted arrow for the postulated flippase mechanism of lipid II inside of the cytoplasm.

Since we postulated a lipid II flippase mechanism the original flippase in the lipid II synthesis MurJ should be considered, which transports this lipid II molecule after synthesis from the cytosol to the extracellular space. Recently, the mechanism of this protein and its conformations were elucidated (41). MurJ belongs to the multidrug/oligosaccharidyl-lipid/polysaccharide (MOP) superfamily, composed of 14 TMHs, which are distinct into the N-lobe (TMH 1-6), the C-lobe (TMH 7-12) and the TMHs 13 and 14, which do not belong to the core transport domain. It is postulated, that a loop between TMH 6 and 7 has an effect on the inner- and outward-facing conformation (42).

The hall mark of BceAB-type transporters is also a extra cellular domain (ECD) of roughly 220 aminoacids, which is localized between TMH 7 and 8. The exact function of this ECD is not known, and intriguingly the sequence homology between the BceAB transporter is rather low (12-15%). As this ECD is very specific for these kind of transporters it likely fulfills an important function, which could be the change of the conformation of the transporter to enable the transport of lipid II from the extracellular space into the cytoplasm.

Conclusively BceAB-type transporter like *Sa*NsrFP are evolutionary conserved in human pathogenic and non-pathogenic strains. Although on sequence level they are less conserved the topology of the protein as well as their encoding operons are conserved. The hallmark is the ECD found on the extra cellular site in between transmembrane helix 7-8. The resistance observed in different BceAB-type transporter studies indicate a common mechanism, which we deduced as flipping of the cell wall precursor lipid II. Some Bce-like operons also encode for additional membrane anchored proteins like NSR of *S. agalactiae* or VraH from *S. aureus* (43), which are very specific for one specific compound. In contrast the BceAB-type ABC-transporter protects the cell against a broad spectrum of antimicrobial peptides with a specific mechanism, demonstrating the high effectiveness.

## Materials and Methods

### Cloning and Expression

Cloning of *nsrFP* from *S. agalactiae* COH1 was performed liked described in Alkhatib et al. (2014) (44) and Reiners et al. (2017) (26) to gain the plasmids pIL-SV *Sa*NsrFP and pIL-SV *Sa*NsrF_H202A_P, latter containing a point mutation in the H-loop which is known to be important for the ATP-hydrolysis (45). These plasmids as well as an empty vector pIL-SVCm were transformed into electrocompetent *L. lactis* NZ9000 cells (46) and the generated strains are termed NZ9000NsrFP, NZ9000NsrF_H202A_P and NZ9000Cm.

The *L. lactis* strains NZ9000NsrFP, NZ9000NsrF_H202A_P were grown in GM17 medium containing 5 μg/ml chloramphenicol. The expression was induced by adding 0.3 nM nisin, cultures were grown at 30°C.

For analyzing the expression, cultures were grown for 5 h and afterwards harvested using a centrifugation step 30 minutes at 5000 x g. The pellets were resuspended with resuspension buffer (50 mM HEPES pH 8.0, 150 mM NaCl, 10 % glycerol) to an OD_600_ of 200 and 1/3 (w/v) 0.5 mm glass beads were added. The cells were lysed and the supernatant was separated by harvesting cell debris as well as glass beads with a 10,000 x g centrifugation step. The membranes were harvested from the supernatant by 100,000 x g centrifugation step. Membrane fractions were prepared with SDS-loading dye (0.2 M Tris–HCl, pH 6.8, 10% (w/v) SDS, 40% (v/v) glycerol, 0.02% (w/v) bromophenol and β-mercaptoethanol). These samples were used for SDS-PAGE and western blot analysis and expressed *Sa*NsrFP proteins detected with a polyclonal antibody against the large extracellular domain of *Sa*NsrP (Davids Biotechnologie, Regensburg, Germany).

### Biological assays

#### Purification of nisin

Nisin was purified with an ion-exchange chromatography as previously described (47) and the concentration determined with RP-HPLC according to Abts et al. (2013) (48).

#### Determination of the half-maximal inhibitory concentration (IC_50_)

The half maximal inhibitory concentration was determined according to Abts et al. (2011) (47). Briefly, *L. lactis* NZ9000Cm, *L. lactis* NZ9000NsrFP and *L. lactis* NZ9000NsrF_H202A_P cells were grown in GM17 medium containing 5 μg/ml chloramphenicol and 0.3 nM nisin at 30°C overnight. Fresh GM17Cm medium with sublethal amount of nisin (0.3 nM) was inoculated with overnight cultures to an OD_600_ of 0.1. A 96-well plate was prepared with a serial dilution of examined antibiotics and subsequent the cell culture was added and plates were incubated at 30°C for 5h. Afterwards the optical density was measured and the IC_50_ values for each strain and antibiotic were calculated (26). To make those values more comparable the fold of resistance was determined by dividing the IC_50_ values of *L. lactis* NZ9000NsrFP and *L. lactis* NZ9000NsrF_H202A_P with the corresponding value for *L. lactis* NZ9000Cm.

#### Growth curve

To detect the growth behavior of the different strains precultures of *L. lactis* NZ9000Cm, *L. lactis* NZ9000NsrFP and *L. lactis* NZ9000NsrF_H202A_P cells were grown in GM17 medium with 5 μg/ml chloramphenicol and 0.3 nM nisin at 30 °C overnight. Freshly prepared GM17Cm medium with 0.3 nM nisin was inoculated with overnight cultures to an OD_600_ of 0.1 and grown to an OD_600_ of 0.4 - 0.5 at 30 °C. These steps were repeated and afterwards the cells were diluted to an OD_600_ of 0.05 in GM17Cm medium containing 0.3 nM nisin. Cells were prepared with additional nisin (1 nM - 40 nM) and bacitracin (0.1 μM - 100 μM) concentrations, respectively. Growth was detected at OD_584_ every 10 minutes with a FLUOstar OPTIMA (BMG Lab technology).

### Cell wall precursor analysis

#### Growth condition and sample preparation

Cells were grown in M17 medium supplemented with 0.5 % glucose and 0.3 nM nisin overnight at 30°C without shaking. At next day 100 ml with 0.5 % glucose and 0.3 nM nisin were inoculated with overnight cultures to on OD_600_ = 0.1. When OD_600_ = 1.2 100 μg/ml bacitracin was added to the cultures to enrich cell wall precursors and cultures were incubated for further 30 min at 30°C. This step was repeated once. (As control a second culture each was harvested before bacitracin was added at an OD_600_=1.2 and cell pellets were stored at −20 °C) After incubation with bacitracin the cells were harvested and the cell pellets were stored at −20 °C. At the next day the cell pellets were resuspended in 25 ml water and cooked for 60 min in boiling water to break the cells. Cell debris were removed by centrifugation (15 min, 500 x g, 4 °C). The supernatant, containing the cell wall precursors, was lyophilisized overnight. Cell pellets were resuspended in 150 μl water and used for LC/MS analysis.

#### LC/MS analysis of cell wall fragments

5 μl of each sample was injected into XCT6330 LC/MSD ultra trap system (Agilent Technologies) equipped with a Nucleosil 100 C18 column (3 μm x 100 mm × 2 mm internal diameter, Dr. Maisch GmbH). The column was used at 40°C. A linear gradient was performed from 0 % up to 10 % eluent B (0.06 % formic acid in acetonitrile) over 25 min with a flow rate of 400 μl/min. The column was re-equilibrated for 10 min with 100 % buffer A (0,1 % formic acid in water). Ionization alternated between positive and negative ion mode with a capillary voltage of 3.5 kV at 350 °C. Extracted ion chromatograms (EIC) in negative ion mode for UDP-MurNAc-Ala-Glu-Lys-Ala-Ala (m/z^-1^ 1148.34 +/-0.1) and UDP-MurNAc-Ala-Glu-Lys-Asp-Ala-Ala (m/z^-1^ 1263.37 +/−0.1) were analyzed with Data Analysis (Bruker), exported as .xy files and presented with GraphPad Prism 6.0.

### Proteom analysis

#### Sample preparation

The *L. lactis* strains NZ9000NsrFP, NZ9000NsrF_H202A_P were grown at 30 °C in GM17 medium containing 5 μg/ml chloramphenicol and 0.3 nM nisin. Precultures were inoculated to an OD_600_ of 0.1 and grown to the exponential growth phase before a main culture was inoculated to an OD_600_ of 0.1. The cells were harvested after 5 h at 5000 x g and the pellet was resuspended in phosphate buffer pH 7 to an OD_600_ of 200 and 1/3 (w/v) 0.5 mm glass beads were added. The cells were lysed and the supernatant was separated by a centrifugation of 10,000 x g.

Protein concentration was determined by means of Pierce 660 nm Protein Assay (Fischer Scientific, Schwerte, Germany) and 10 μg protein per sample were loaded on an SDS-PAGE for in-gel-digestion. The isolated gel pieces were reduced, alkylated and underwent afterwards tryptic digestion. The peptides were resolved in 0.1 % trifluoracetic acid and subjected to liquid chromatography.

#### LC-MS analysis

For the LC-MS analysis a QExactive plus (Thermo Scientific, Bremen, Germany) connected with an Ultimate 3000 Rapid Separation liquid chromatography system (Dionex / Thermo Scientific, Idstein, Germany) equipped with an Acclaim PepMap 100 C18 column (75 μm inner diameter, 25 cm length, 2 mm particle size from Thermo Scientific, Bremen, Germany) was applied. The length of the LC gradient was 120 minutes. The mass spectrometer was operating in positive mode and coupled with a nano electrospray ionization source. Capillary temperature was set to 250°C and source voltage to 1.4 kV. In the QExactive plus mass spectrometer for the survey scans a mass range from 200 to 2000 m/z at a resolution of 70,000 was used. The automatic gain control was set to 3,000,000 and the maximum fill time was 50 ms. The 10 most intensive peptide ions were isolated and fragmented by high-energy collision dissociation (HCD).

#### Computational mass spectrometric data analysis

Proteome Discoverer (version 2.1.0.81, Thermo Fisher Scientific, Bremen, Germany) was applied for peptide/protein identification applying Mascot (version 2.4, Matrix Science, London, UK) as search engine employing the EnsemblBacteria database (*Lactococcus lactis* subsp. Cremoris NZ900; date 03-11-2019). A false discovery rate of 1% (p ≤ 0.01) on peptide level was set as identification threshold. Proteins were quantified with Progenesis QI for Proteomics (Version 2.0, Nonlinear Dynamics, Waters Corporation, Newcastle upon Tyne, UK). Only proteins containing at least two unique peptides were taken into consideration. For the calculation of enriched proteins in the groups a 5 % false discovery rate and a minimum fold change of two was used.

The mass spectrometry proteomics data has been deposited to the ProteomeXchange Consortium via the PRIDE partner repository with the data set identifier PXD017318.

## Acknowledgments

The authors are thankful to Jelle Postma and Stefanie Weidtkamp-Peters from the Center for Advanced Imaging, Heinrich-Heine-University Germany, for valuable suggestions and great support during fluorophore studies. This work was funded by the Deutsche Forschungsgemeinschaft (DFG, German Research Foundation) – 270650915/GRK 2158.

## Supporting Information

**SI Figure 1:**
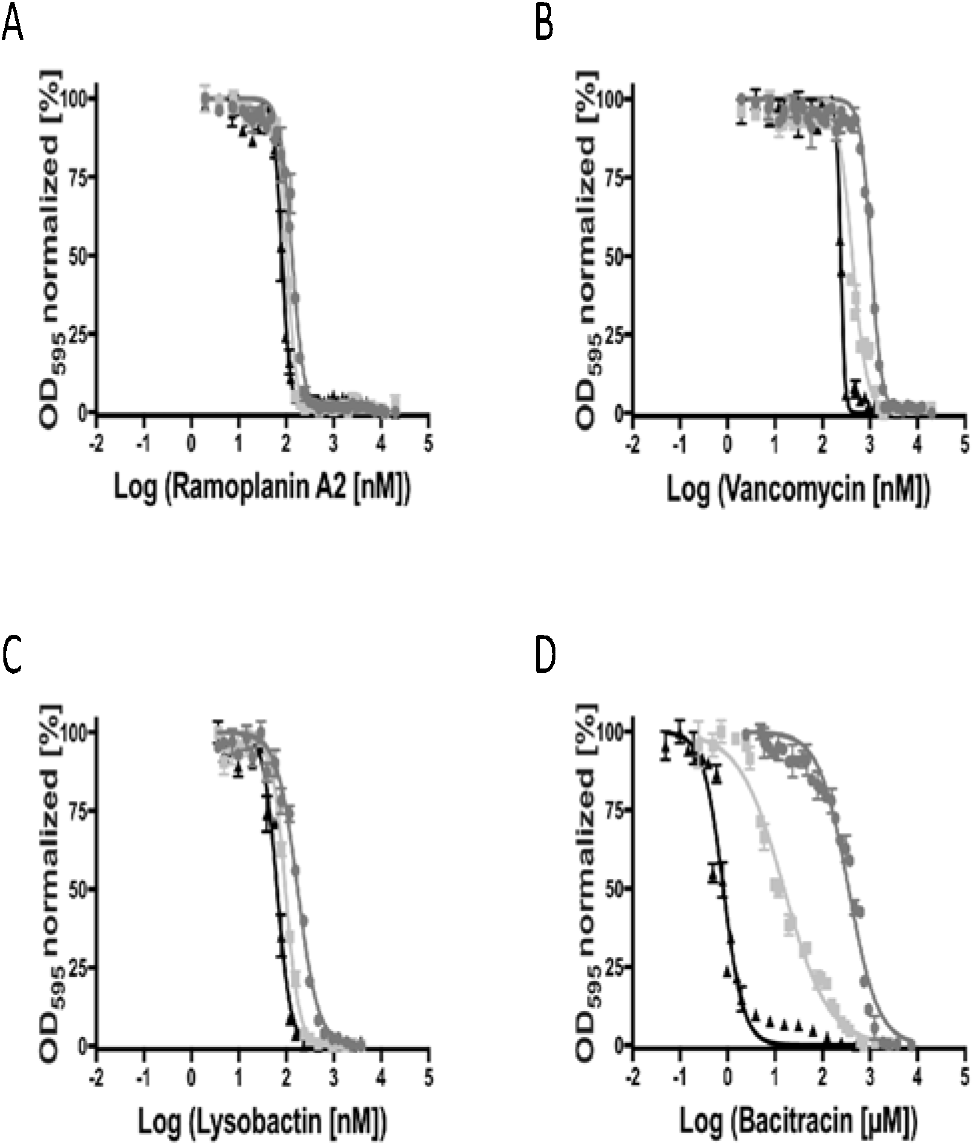
**Representative inhibitional growth curves** of **A) Ramoplanin A2, B) Vancomycin, C) Lysobactin** and **D) Bacitracin.** Normalized OD_595_ is plotted against the logarithmic concentration of the antibiotic. NZ9000Cm is demonstrated in black, NZ9000NsrF_H202A_P in light grey and NZ9000NsrFP in grey.

**SI Table 1:** **Measured IC_50_ values and calculated fold of resistance** for the antibiotics Ramoplanin A2, Vancomycin, Lysobactin and Bacitracin and for the strains NZ9000Cm, NZ9000NsrF_H202A_P and NZ9000NsrFP.

| Antibiotic | <u>*L. lactis* NZ9000Cm</u> | <u>*L. lactis* NZ9000NsrFP</u> |  | <u>*L. lactis* NZ9000NsrF<sub>H202A</sub>P</u> |  |
| --- | --- | --- | --- | --- | --- |
|  | IC<sub>50</sub> [nM] | IC<sub>50</sub> [nM] | Fold of resistance | IC<sub>50</sub> [nM] | Fold of resistance |
| Ramoplanin A2 | 73.00 ± 18.2 | 120.52 ± 34.2 | 1.65 ± 0.5 | 91.67 ± 20.5 | 1.26 ± 0.3 |
| Vancomycin | 213.57 ± 27.0 | 1077.92 ± 263.5 | 5.05 ± 1.2 | 325.38 ± 117.1 | 1.52 ± 0.5 |
| Lysobactin | 30.77 ± 16.8 | 181.68 ± 36.5 | 5.91 ± 1.2 | 101.80 ± 52.2 | 3.31 ± 1.8 |
| Bacitracin | 937.64 ± 93.9 | 327500.00 ± 60883.9 | 349.15 ± 64.9 | 12854.8 ± 8517.2 | 13.70 ± 9.1 |

**SI Figure 2:**
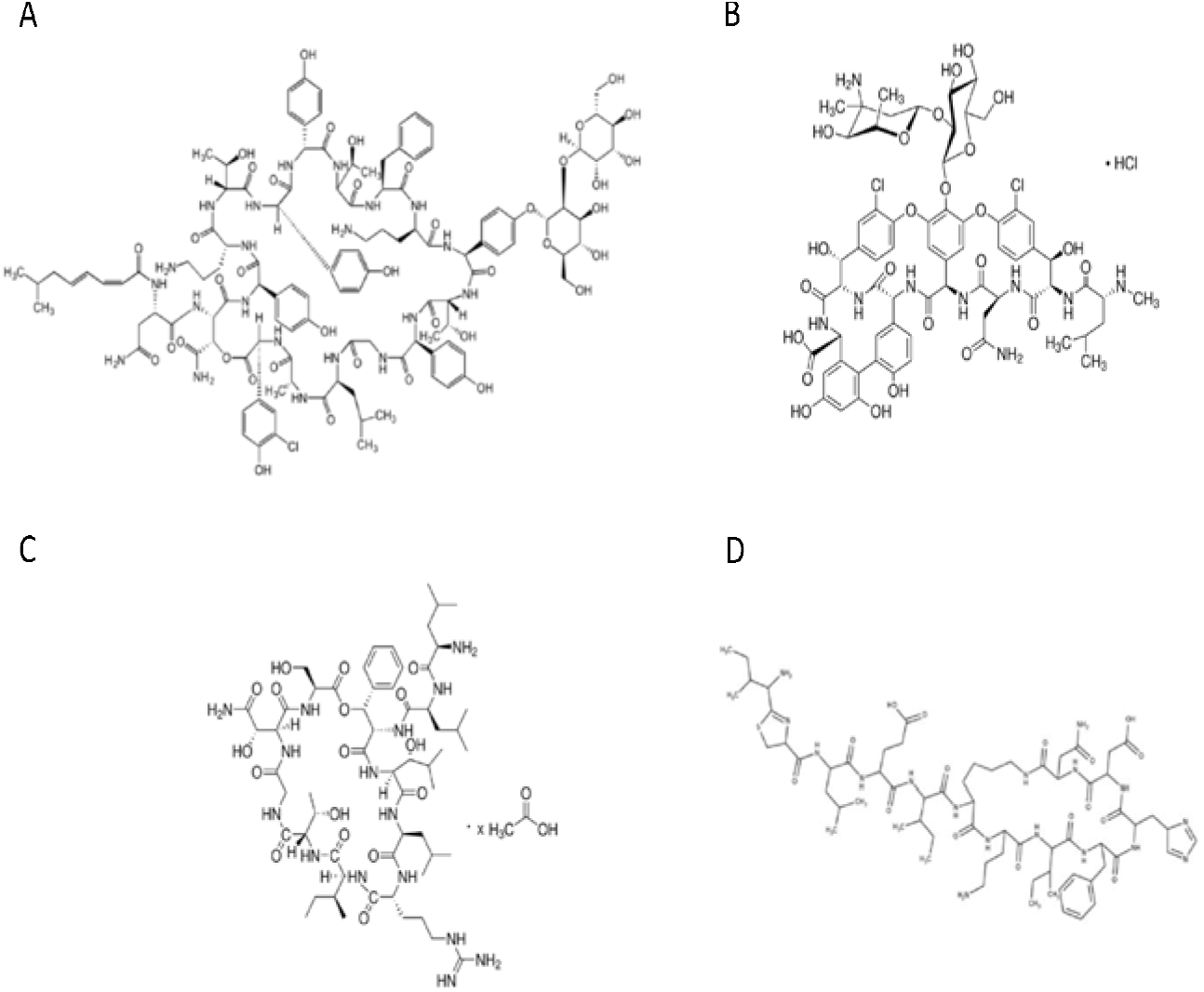
**Structures of A) Ramoplanin A2** from AdipoGen life sciences, **B) Vancomycin** from Fluka Analytical, **C) Lysobactin** from Sigma life sciences and **D) Bacitracin** from Fisher BioReagents.

